# *Plasmodium falciparum* genetic diversity in coincident human and mosquito hosts

**DOI:** 10.1101/2022.05.05.490756

**Authors:** Zena Lapp, Andrew A Obala, Lucy Abel, David A Rasmussen, Kelsey M Sumner, Elizabeth Freedman, Steve M Taylor, Wendy Prudhomme-O’Meara

## Abstract

Population genetic diversity of *P. falciparum* antigenic loci is high despite large bottlenecks in population size during the parasite life cycle. The extent of this diversity in human blood-stage infections, following expansion from a small number of liver-stage schizonts, has been well described. However, little is known about parasite genetic diversity in the vector, where a similar bottleneck and expansion occurs following parasite mating and where parasite genotypes from several different human infections may accumulate. We assessed parasite genetic diversity within human and mosquito *P. falciparum* infections collected from the same households during a 14-month longitudinal cohort study using amplicon deep sequencing of two antigenic gene fragments (*ama1* and *csp)*. To a prior set of infected humans (n=1175/2813; 86.2% sequencing success) and mosquito abdomens (n=199/1448; 95.5% sequencing success), we added sequences from infected mosquito heads (n=134/1448; 98.5% sequencing success). Across all sample types we observed 456 *ama1* and 289 *csp* unique haplotypes. While both hosts contained many rare haplotypes, population genetic metrics indicated that the overall and sample-level parasite populations were more diverse in mosquitoes than in humans, and infections were more likely to harbor a dominant haplotype in humans than in mosquitoes (based on relative read abundance). Finally, within a given mosquito there was little overlap in genetic composition of abdomen and head infections, suggesting that infections may be cleared from the abdomen during a mosquito’s lifespan. Taken together, our observations provide evidence for the role of the mosquito vector in maintaining sequence diversity of malaria parasite populations.

**Significance statement:** Concurrent infections with multiple strains of *Plasmodium falciparum*, the leading causative agent of death due to malaria, are common in highly endemic regions. During transitions within and between the parasite’s mosquito and human hosts, population bottlenecks occur, and distinct parasite strains may have differential fitness in the various environments encountered. These bottlenecks and fitness differences may lead to differences in strain prevalence and diversity between hosts. We investigated differences in genetic diversity between *P. falciparum* parasites in human and mosquito hosts and found that, compared to human parasite populations and infections, mosquito populations and infections were more diverse. This suggests that the mosquito vector may play a role in in maintaining sequence diversity in malaria parasite populations.

## Introduction

*Plasmodium falciparum* has a complex life cycle that requires it to navigate multiple cellular and host transitions to sustain transmission. These include transitions both between human and mosquito hosts and between compartments within those hosts. In addition, distinct genotypes may be co-transmitted between hosts in a single bite or may accumulate within a host owing to serial super-infections. Such infections consisting of many different strains are particularly commonplace in highly endemic settings such as some regions of sub-Saharan Africa (1), promoting both outcrossing in mosquito hosts and competition in human hosts. These factors, coupled with population bottlenecks and selective pressures encountered by *P. falciparum* throughout its life cycle, shape overall patterns of parasite genetic diversity (2).

Comparative population genetics of *P. falciparum* between the hosts and cellular environments through which the parasite transitions in natural cycles of transmission remains relatively unexplored. Several studies have compared markers of drug-resistance loci between hosts, and an early report from Zambia observed very different allele frequencies in humans and mosquitoes (3, 4), suggesting differences in parasite population structure between hosts. However, subsequent reports from other settings using different genetic markers have not consistently observed this phenomenon (5, 6). As these studies used marker genes with few polymorphisms, analyses of individuals with complex co-infections was limited. While microsatellite markers overcome some of these limitations (7, 8), prior studies have not to our knowledge contrasted the genetic composition or diversity of highly polymorphic targets in naturally-occurring infections of humans and mosquitoes that are participating in co-incident transmission networks. By exploring these phenomena more closely, we can better understand what factors contribute to the diversity of malaria parasite populations.

We investigated variability in *P. falciparum* genetic diversity across human and mosquito hosts in a highly endemic area of Western Kenya. During a 14-month longitudinal cohort study, we detected *P. falciparum* parasites in human participants and in the heads and abdomens of resting Anopheline mosquitoes collected from their households (1). From each *P. falciparum* infection, we used amplicon deep sequencing of polymorphic segments of the parasite genes encoding apical membrane antigen 1 *(ama1)* and circumsporozoite protein *(csp)* to catalog complex *P. falciparum* infections in human blood, mosquito abdomens, and mosquito heads. We previously reported that parasite multiplicity of infection (MOI) as expressed by either marker was higher in mosquito abdomens harboring recently-ingested parasites than humans harboring blood-stage parasites (1). Here, we examine more carefully the differences between host compartments in haplotype diversity and relative abundance both within a given host and at the population level. Based on our previous observation, as well as the robust immune defenses against *P. falciparum* in humans (9), we hypothesized that the mosquito *P. falciparum* haplotype population would be more diverse than that of humans.

## Results

### Data overview and analytic population

Samples were collected over the course of 14 months (June 2017 - July 2018) from 38 households in three Kenyan villages. Mosquitoes were aspirated weekly from each household and blood samples from household members were collected monthly. To the previously reported data on humans and mosquito abdomens (1), we added data from mosquito heads. Over a third of human samples (41.8%; 1175/2813) contained *P. falciparum*, compared to 13.7% (199/1448) of mosquito abdomens and 9.2% (134/1462) of mosquito heads (**Figure S1**). Of these, sequencing of at least one marker was successful in 86.2% (1013/1175) of human, 95.5% (190/199) of mosquito abdomen, and 98.5% (132/134) of mosquito head infections. Haplotype information from these 1013 infections in 224 people and 322 infections in 244 mosquitoes constituted the analytic population.

### Mosquito head infections are not a subset of their abdomen infections

Parasite development within the mosquito host begins in the abdomen following which sporozoites must traverse the midgut wall to reach the salivary glands in the head; however, it is not known how quick and comprehensive is this egress. We hypothesized that if both midgut and salivary gland infections persist throughout the mosquito’s lifespan (i.e. incomplete egress from the midgut), haplotypes in a mosquito’s head would be a subset of those in the abdomen. Among mosquitoes in which at least one compartment was infected, *P. falciparum* was detected in both the abdomen and the head in 89/238 (37.4%), in only the abdomen in 108/238 (45.4%), and in only the head in 41/238 (17.2%) (**Figure 1A**). The latter finding suggests that infections may be completely cleared from the abdomen within the span of a mosquito’s lifetime.

**Figure 1:**
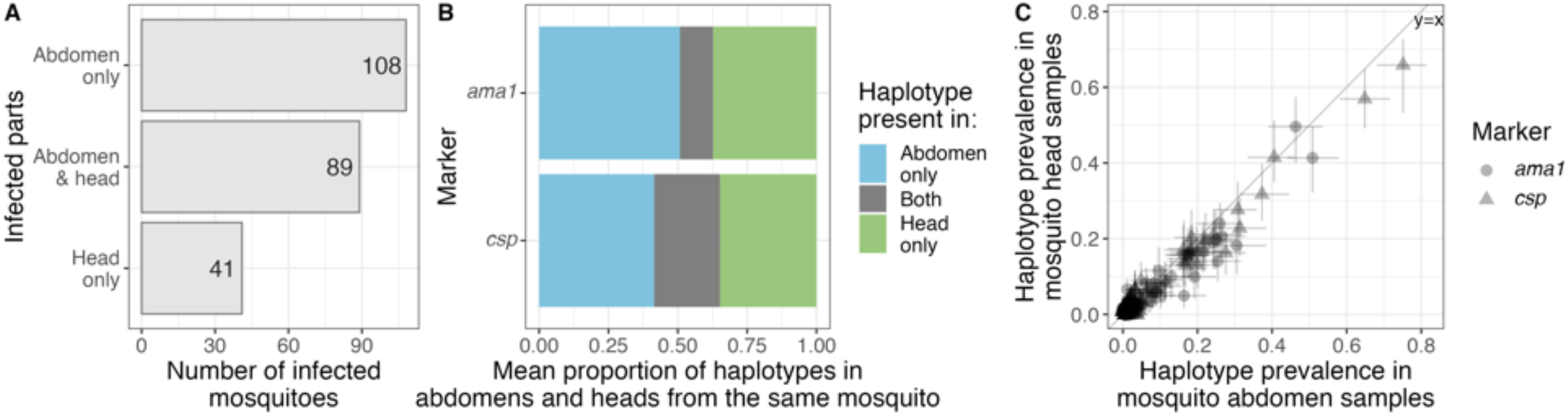
Mosquito abdomens and heads do not contain similar infections. (A) *P. falciparum* infection of mosquito abdomens and heads of mosquitoes for which both compartments were tested using PCR. (B) For 89 mosquitoes with infections of both the abdomen and the head, the proportion of the set of haplotypes in the mosquito found in the abdomen only, head only, or both. The mean counts for each of the three groups were used to obtain the proportions. (C) Prevalence of each *ama1* and *csp* haplotype in the mosquito abdomen population compared to the mosquito head population. Each dot represents a unique *ama1* or *csp* haplotype, and bars indicate the 95% bootstrapped confidence intervals.

We next compared the haplotype compositions of infections in the 89 mosquitoes in which *P. falciparum* was detected in both the head and the abdomen. We calculated the percentage of *ama1* or *csp* haplotypes found only in the head or the abdomen, or observed in both compartments (i.e. the Jaccard distance; intersect/union) within each mosquito. While some haplotypes were observed in both compartments of a given mosquito (mean for *ama1:* 12.0%, *csp*: 23.7%), the majority of haplotypes were either private to the abdomen (mean for *ama1:* 50.7%, *csp*: 41.5%) or head (mean for *ama1:* 37.3%, *csp*: 34.8%) (**Figures 1B, S2A-B**). Despite this limited overlap, sharing between abdomens and heads from the same mosquito was higher than sharing between random pairs of abdomens and heads (**Figure S2C**; Kolmogorov-Smirnov p ≤ 1e-10 for both markers).

To determine whether the differences in haplotype composition between abdomen and head infections within a single mosquito corresponded to differences at the host population level, we compared between mosquito compartments haplotype population-level prevalences, defined as the number of samples in which a haplotype was observed. Both *ama1* and *csp* haplotype prevalences were similar between mosquito abdomen and head populations (**Figure 1C**), suggesting that the transition from oocyst to sporozoite does not alter the diversity of circulating parasites. Owing to this population-level similarity in prevalences and our observation that abdomen and head parasite populations from the same mosquito appear to frequently represent different infections, we subsequently performed all comparisons between the two *P. falciparum* hosts: humans and mosquitoes, where mosquito samples included both abdomen and head samples.

### The *P. falciparum* population in mosquitoes is more diverse than the population in humans

To investigate signatures of differential bottlenecks or selection during parasite transition between mosquito and human hosts, we compared population-level differences in parasite haplotype prevalence among mosquitoes and humans, where differences in prevalence may indicate differential bottlenecks or selection. Across all infections, we observed high haplotype richness, with 456 *ama1* and 298 *csp* distinct haplotypes. The vast majority of these were low-frequency haplotypes, many of which were observed in only one host (**Figures S3-4**). Among 54 distinct haplotypes (both *ama1* and *csp*) with a prevalence above 5% across all samples, we observed 28 haplotypes with differential prevalence across hosts: 19 more common in mosquito infections and 9 in human infections (**Figure 2A**), consistent with our observation of higher average mosquito MOIs (1) (**Figure S5**).

**Figure 2:**
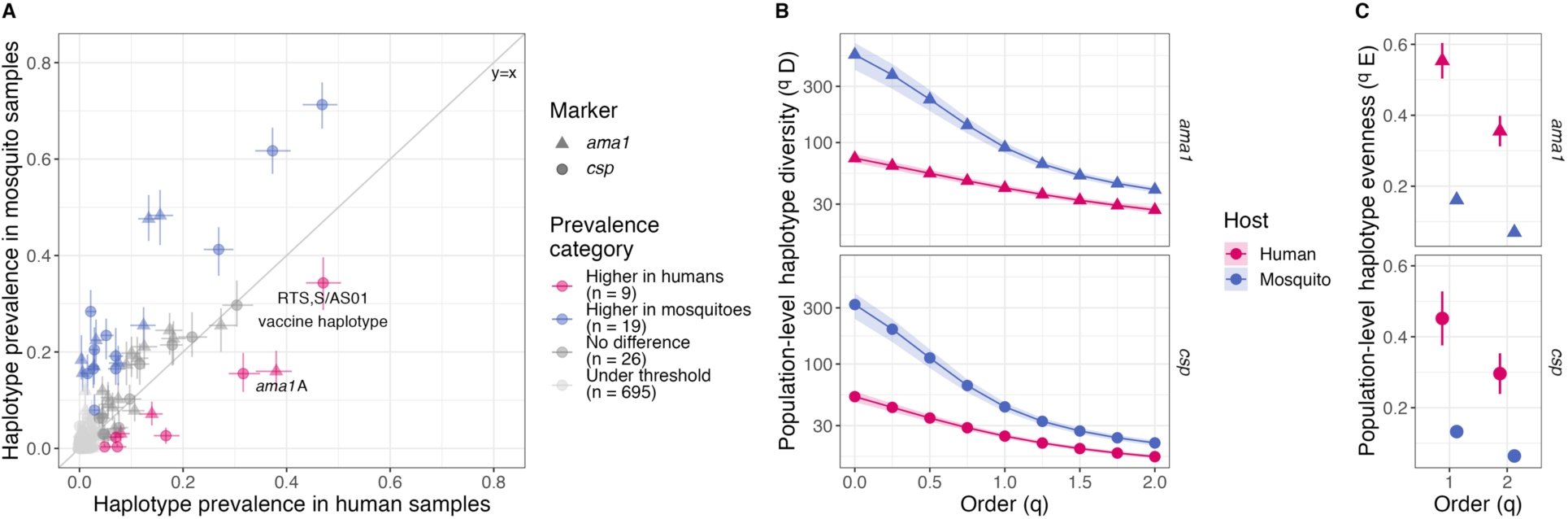
The mosquito *P. falciparum* population is more diverse than the human *P. falciparum* population. (A) Prevalence of each *ama1* and *csp* haplotype in the human population compared to the mosquito population. Each dot represents a unique *ama1* or *csp* haplotype, and bars indicate the 95% bootstrapped confidence intervals. The lower threshold was defined as haplotypes observed in fewer than 5% of combined human and mosquito samples. Haplotype prevalences were considered higher in one compartment if the 95% bootstrapped confidence intervals didn’t overlap the expected prevalence (i.e. the overall prevalence across all samples). (B) Diversity of *P. falciparum* populations by host and genetic marker across orders of diversity. Ribbons are bootstrapped 95% confidence intervals. Higher values indicate more diversity. The slope of the line across orders *q* is a measure of haplotype evenness in the population. (C) Haplotype evenness of human and mosquito samples. Bars are bootstrapped 95% confidence intervals. Higher values indicate more similar prevalence of haplotypes in the population.

We next used haplotype prevalence to quantify population-level diversity across orders of diversity *(q)* ranging from equal weight to each haplotype *(q* = 0, equivalent to haplotype richness or the number of distinct haplotypes observed) to downweighting rare haplotypes (q = 2, effective number of highly abundant haplotypes) (10). The mosquito parasite population was more diverse than the parasite population in human hosts (**Figure 2B**; **S6A**). This trend is consistent even when accounting for multiple samples per host, different sampling schemes between hosts, differences in MOI, haplotypes with rare variants, and limitations of using empirical diversity (**Figure S6B**). Moreover, as evidenced by the steeper decline in diversity with increasing *q* in mosquitoes relative to humans (**Figures 2B**), the mosquito parasite population contained more uneven haplotype prevalences than the human parasite population (**Figure 2C**), indicating that mosquitoes contained a larger relative number of infrequent haplotypes. Even so, higher diversity in the mosquito host is still apparent when downweighing the contribution of these minor haplotypes. Taken together, these results indicate that there may be a greater relative loss in diversity across the transition from mosquitoes to humans than humans to mosquitoes.

### Dominant haplotypes within infections are more common in humans than mosquitoes

In addition to lower population-level diversity in humans compared to mosquitoes, we also observed lower within-sample diversity (**Figure S5**) and proportionately more monoclonal infections in humans (**Figure 3A**; both Fisher’s exact p < 1e-12). To further investigate whether human infections are more often dominated by one or a few haplotypes relative to mosquito infections, we calculated for each infection the haplotype evenness, which examines the haplotype abundance within an infection based on sequencing reads. A lower value indicates that the infection consists of mostly reads from a single haplotype or, in other words, is dominated by a majority haplotype. Median evenness values for mosquito infections were higher *(ama1:* 0.88; *csp:* 0.84) than those in human infections *(ama1:* 0.51; *csp:* 0.67) (all p < 1e-4) (**Figure 3B**). This observation was robust to taking the maximum evenness among the two markers and to differences in haplotype filtering (all p < 1e-10; **Figure S7**). This differential composition of polyclonal *P. falciparum* infections between hosts supports a differential in selective landscapes that may further enable the preservation of diverse *P. falciparum* populations in Anopheline mosquitoes.

**Figure 3:**
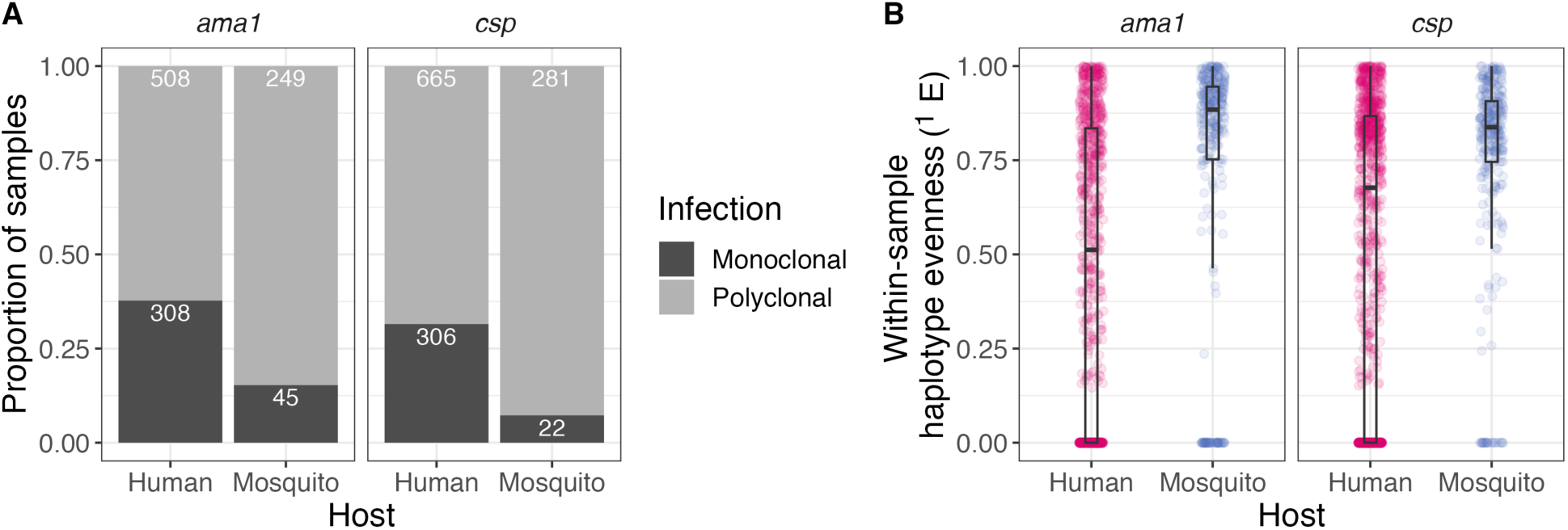
Compared to mosquito samples, human samples are more often dominated by a single haplotype. (A) Proportion of samples with monoclonal and polyclonal infections. Numbers are counts for each category. (B) Distributions of within-sample evenness (q = 1) by genetic marker and host. Lower values indicate more dominance by individual haplotypes within the strain mixture.

## Discussion

We compared *P. falciparum* genetic diversity across several host compartments that the parasite must successfully navigate to sustain transmission. Parasite genetic diversity was increased relative to humans during the mosquito stages, although this incremental diversity in mosquitoes appears to be transmitted only infrequently to humans. In addition, individual infections were composed differently in mosquitoes and humans, with human infections more commonly harboring dominant members. Collectively, our observations suggest that mosquito-stage infections participate the maintenance of diversity in *P. falciparum* parasite populations not only through recombination, but also by acting as a reservoir of sequence diversity.

We observed, using multiple metrics, more parasite genetic diversity in mosquitoes compared to humans. This high diversity contrasts with the known marked reduction in parasite biomass during the transition from the human to the mosquito abdomen (11), which might be expected to constrain parasite diversity. One potential explanation for this is the possibility of cryptic genotypes in humans undetected by marker sequencing; this has been reported in experimental studies (7), though the large range of MOIs we observed in humans suggests that these infections were not systematically undersampled. Alternatively, the reduced diversity in humans could result from large reductions in population size and negative selective pressures as the parasite passes from mosquitoes, through the human liver, and into the blood stage. Mosquitoes are the location of parasite sexual recombination and therefore certainly provide a site for genomic diversification, but this seems unsuited to explain the diversity of these short segments in *ama1* and *csp* that do not harbor known recombination hotspots (12). A probable contributor to this high mosquito diversity is multiple or interrupted feeds on infected hosts, which would allow strains to accumulate in the mosquito abdomen. This feeding behavior has been reported for *Anopheles gambiae* and may be enhanced by human *P. falciparum* infection (13, 14). Additionally, *P. falciparum* adaptation to evade the immune system of local *Anopheles* strains (15), as well as imposition of selective pressures on the *Anopheles* vector to the parasites’ benefit (16), may reduce differences in fitness between distinct parasite strains within the mosquito and lead to an accumulation of genetic diversity at the population level. However, some of these novel strains may be unfit to survive the human host. Indeed, prior work on arbovirus infection found an accumulation of mutations in mosquitoes that led to fitness costs during vertebrate infection (17). Despite these plausible explanations for constrained diversity in humans and higher diversity in mosquitoes, the mechanism by which mosquitoes maintain such high parasite diversity when their parasite population is necessarily sampled from the less diverse human population remains to be fully elucidated.

Within individual infections, we observed higher dominance of haplotypes in human compared to mosquito infections, while on a larger scale, the *P. falciparum* haplotype population was more evenly distributed among humans than among mosquitoes. These differences may result from the differential selection landscapes between hosts, in particular for the proteins encoded by our gene targets, AMA1 and CSP, which harbor epitopes that are known targets of functional human immunity (18). In humans, the concurrent maintenance in the population of multiple viable alleles due to balancing selection, paired with the removal of deleterious alleles due to negative selection, could produce a relatively high evenness of haplotypes in the human parasite population even as individual infections are shaped by directional selection resulting from individual host immune responses. In contrast, the relative lack of differential fitness in the mosquito host described above may lead to even parasite strain abundances within a mosquito.

Comparison of paired abdomens and heads from the same mosquito revealed striking differences between *P. falciparum* presence and haplotype composition. As expected given the delay between midgut and salivary gland infections, many mosquitoes had haplotypes private to the abdomen that were not present in the head. More surprising was the observation of mosquitoes with haplotypes private to the head that were absent from the abdomen, suggesting that infections do not reliably persist in a mosquito’s abdomen throughout its lifespan. While these differences may again be due to cryptic haplotypes, the identification of mosquitoes with infections in the head but not the abdomen using sensitive PCR detection methods (19, 20) indicates that cryptic haplotypes likely cannot explain all of the observed differences. Despite these discrepancies between abdomens and heads from a given mosquito, at the population level haplotype composition and diversity were similar between mosquito abdomens and heads, suggesting that the selective pressures for or against certain haplotypes (or lack thereof) may be similar in these two compartments.

Our findings highlight the role of the mosquito host in influencing the sequence diversification of *P. falciparum* parasites. A unique feature of genetic diversity in *P. falciparum* compared to other organisms is the preponderance of low-frequency alleles (21). A prior modeling study suggested that this phenomenon may be the result of the complex, “unconventional” life cycle of *P. falciparum*, specifically the bottlenecks and host transitions that intensify both random genetic drift as well as natural selection (2). Consistent with this, we observed many haplotypes private to one host, which was more prominent in mosquitoes. As noted above, meiotic recombination is unlikely to be the main contributor to the diversity we cataloged, and the mechanisms by which these low-frequency and private alleles arise remains obscure. However, our observations furnish compelling evidence for a role of the mosquito vector in accumulating genetic diversity in genic regions likely not under positive selection in mosquitoes. While this diversity appears to be selected against in humans, it nevertheless acts as a continual supply of novel alleles and allelic combinations that may be exploited by the parasite during human infection.

This study has limitations. First, the inability to sample parasites from mosquitoes without sacrificing them precludes a comprehensive study of paired mosquito abdomen and head infections over time. Even so, we were still able to identify similarities and differences between the haplotype populations in these two compartments. Additionally, the mosquito and human sampling schemes were different, potentially biasing sampling comprehensiveness between hosts. To mitigate the risk that this potential imbalance influenced our results, we performed comparative population analyses using empirical methods with a fixed coverage threshold (10) and sensitivity analyses. Finally, many of the human and mosquito infections had very low parasite densities, which not only increases the possibility of failing to detect infections, but also increases the possibility of false haplotype discovery (22). To reduce the inclusion of false haplotypes to the greatest extent possible, we performed strict haplotype censoring to remove potential false positives (1) and performed sensitivity analyses on key findings to determine whether haplotype filtering criteria influenced the results.

In conclusion, our comparison of *P. falciparum* haplotypes observed in natural, coincident infections of humans, mosquito abdomens, and mosquito heads revealed greater genetic diversity in mosquito than human populations and infections. This provides evidence for the role of the mosquito vector in maintaining the sequence diversity of malaria parasite population.

## Materials and methods

### Ethics statement

All adults, and parents or legal guardians for individuals under 18 years old, provided written informed consent. Children over 8 years old also provided verbal assent. The study was approved by the ethical review boards of Moi University (2017/36) and Duke University (Pro00082000).

### Study design and sampling

The study design and sample processing have been described previously (1). Briefly, a longitudinal cohort of participants (1 year of age or older) residing in 38 households in three villages in Western Kenya were followed from June 2017 to July 2018. For each participant, dried blood spots (DBS) were collected monthly and any time participants had malaria symptoms. One morning each week, indoor resting mosquitoes were collected from participant households using vacuum aspiration, and following morphologic identification, the abdomen was separated from the head and thorax of female *Anopheles* mosquitoes. Genomic DNA was isolated from DBS, mosquito abdomens, and mosquito heads; *P. falciparum* was detected in these extracts using a real-time PCR assay. Segments of approximately 300 nucleotides of the *P. falciparum ama1* and *csp* genes were amplified, sequenced on an Illumina MiSeq platform, and haplotype inference was performed using DADA2 v1.8 (23) with custom read- and haplotype-filtering. The output was a set of quality-filtered *ama1* and *csp* reads and corresponding parasite haplotypes for each *P. falciparum* infection.

We performed parallel analyses of amplicon deep-sequenced segments of the *P. falciparum ama1* and *csp* marker genes. Since *ama1* and *csp* are unlinked markers found on different chromosomes, to some extent these parallel analyses can be considered pseudo-replicates, where similar results for both markers increases confidence in our findings.

### Within-mosquito comparison

For each mosquito with a *P. falciparum* infection in both the abdomen and the head, the Jaccard distance (24) was calculated for the haplotypes in the abdomen-head pair:

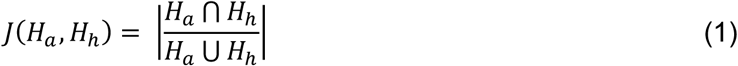

Where *H*_***a***_ is the set of haplotypes in the abdomen and *H*_***h***_ is the set of haplotypes in the head.

### Haplotype prevalence

For each haplotype, population-level prevalence was determined for 5 distinct populations: the entire sample set, all human samples, all mosquito samples, mosquito abdomens, and mosquito heads. Prevalence was calculated as the proportion of samples harboring that haplotype. 95% confidence intervals were computed from 100 bootstrapped datasets. Haplotypes were considered low-frequency if they occurred in fewer than 5% of all samples. Haplotypes that were not low-frequency (i.e. above a threshold of 5% prevalence) were considered higher in a given compartment if the range of bootstrapped prevalences did not overlap the expected prevalence (i.e. the overall prevalence across all samples).

### Randomized minimum spanning trees

To visualize the relatedness of haplotypes, we calculated pairwise distances using the dist.dna() function in the R package ape v5.6-2 (25) with the K80 evolutionary model, computed randomized minimum spanning trees (26) using the rmst() function in pegas v1.1 (27), and visualized the trees in ggtree v3.0.4 (28).

### Diversity and evenness

For analyses between humans and mosquitoes, all mosquito abdomen and head samples were considered mosquito samples, providing a maximum of 2 samples from each mosquito.

#### Population-level

For the set of mosquito samples and the set of human samples, we calculated the population-level diversity of haplotypes, rarefaction curves, and population evenness using the R packages iNEXT.4steps v1.0.1 (10) and iNEXT.3D v1.0.1 (29).

Diversity was calculated using the following equation (10, 30):

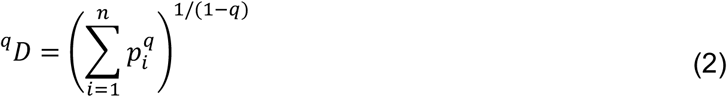

Where *q* is the order of diversity, *n* is the number of distinct haplotypes, and *p*, is the prevalence of haplotype *i* in the sample set. D was computed across a range of values *q* between 0 and 2, where higher numbers correspond to upweighting haplotypes that are more abundant in the overall population. True diversity was not accurately calculable for low orders of diversity (*q* < 1) due to an abundance of unsampled rare haplotypes. Therefore, to enable comparison of diversity between the human and mosquito haplotype populations, we calculated the empirical diversity at a standardized coverage of the host population’s haplotypes (90.1% for *ama1* and 94.5% for *csp)*.

Sensitivity analyses were performed using (1) true (asymptotic) diversity (for *q* ≥ 1), (2) a subsampled dataset including the same number of host samples per week (to account for differences in mosquito and human sampling schemes), (3) a subsampled dataset including only one sample per host (to account for multiply sampled hosts), (4) by defining *p*, as the frequency of haplotypes in the total set of haplotypes (to account for differences in MOI between infections), and (5) a dataset including only haplotypes with variation at amino acid positions that are variable in both hosts (to limit potential false positives by using this stricter set of haplotype filtering criteria).

We calculated haplotype evenness using the following equation (31):

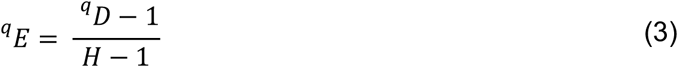

Where *H* is the haplotype richness, or the number of distinct haplotypes in the population. For *q* = 0 evenness is defined as 1, and for *H* = 1, evenness is defined as 0.

#### Within-sample

For each sample, we computed haplotype diversity and evenness using equations 1 and 2. In this case, *p*_***i***_ in equation 1 is the relative read abundance of each haplotype, ^***q***^ *D* is the within-host diversity, and *H* is the MOI of the infection.

To compare evenness between human and mosquito hosts, we computed a zero-one inflated Beta regression model using the R package gamlss v5.4-1 (32) with host as the main exposure, evenness as the outcome, log2-transformed haplotype reads as a covariate, and individual as a random effect. To determine whether incorporating information from both markers influenced differences in evenness between hosts, for each sample we selected the highest evenness value (between *ama1* and *csp)* and compared these values between humans and mosquitoes. Finally, to explore if evenness values were biased by the initial enforcement of haplotype quality-filtering criteria that were partially based on within-sample haplotype proportion, we performed a sensitivity analysis using unfiltered haplotypes. These haplotypes were inferred by DADA2 v1.8 (23) from input reads which passed upstream read quality-filtering. Using these unfiltered haplotypes, we used the same methods as above to compute and compare evenness.

### Data analysis and visualization

Comparison across groups was performed using Wilcoxon rank-sum tests, Fisher’s exact tests, or Kolmogorov-Smirnov tests. All data analysis and visualization was performed in R v4.1.1 (33) and RStudio v2021.9.0.351 (34) using the following packages: tidyverse v1.3.1 (35), ape v5.6-2 (25), cowplot v1.1.1 (36), scales v1.1.1 (37), and ggtext v0.1.1 (38). All data and code to reproduce the analyses and figures can be found on GitHub (https://github.com/duke-malaria-collaboratory/parasite-host-comparison).

## Acknowledgements

We thank the field technicians in Webuye for their engagement with the study participants: Khaoya, L. Marango, E. Mukeli, E. Nalianya, J. Namae, L. Nukewa, E. Wamalwa, and A. Wekesa; J. Kipkoech Kirui (AMPATH) for operational assistance; V. Liao, A. Nantume, S. Kim, and J. Saelens (each of Duke University) for laboratory sample and data processing; and C. Markwalter PhD (Duke University) for thoughtful discussion. Ultimately, we are indebted to the households for their participation in this study. Preliminary results were presented at the Annual Meeting of the American Society of Tropical Medicine and Hygiene in November 2020.

## Author contributions

DAR, KMS, SMT, WPO, and ZL conceptualized the study. EF, SMT, WPO, KMS, and ZL developed the methodology. ZL and KMS curated the data, performed formal analysis, and performed the computer programming. ZL visualized the data. WPO, SMT, KMS, and DAR acquired funding. ZL, LA, EF, and KMS performed the investigation. SMT, WPO, AO, and LA administered and supervised the project. ZL, SMT, and WPO wrote the original manuscript draft. All authors reviewed and edited the manuscript.

## Supplementary figures

**Figure S1:**
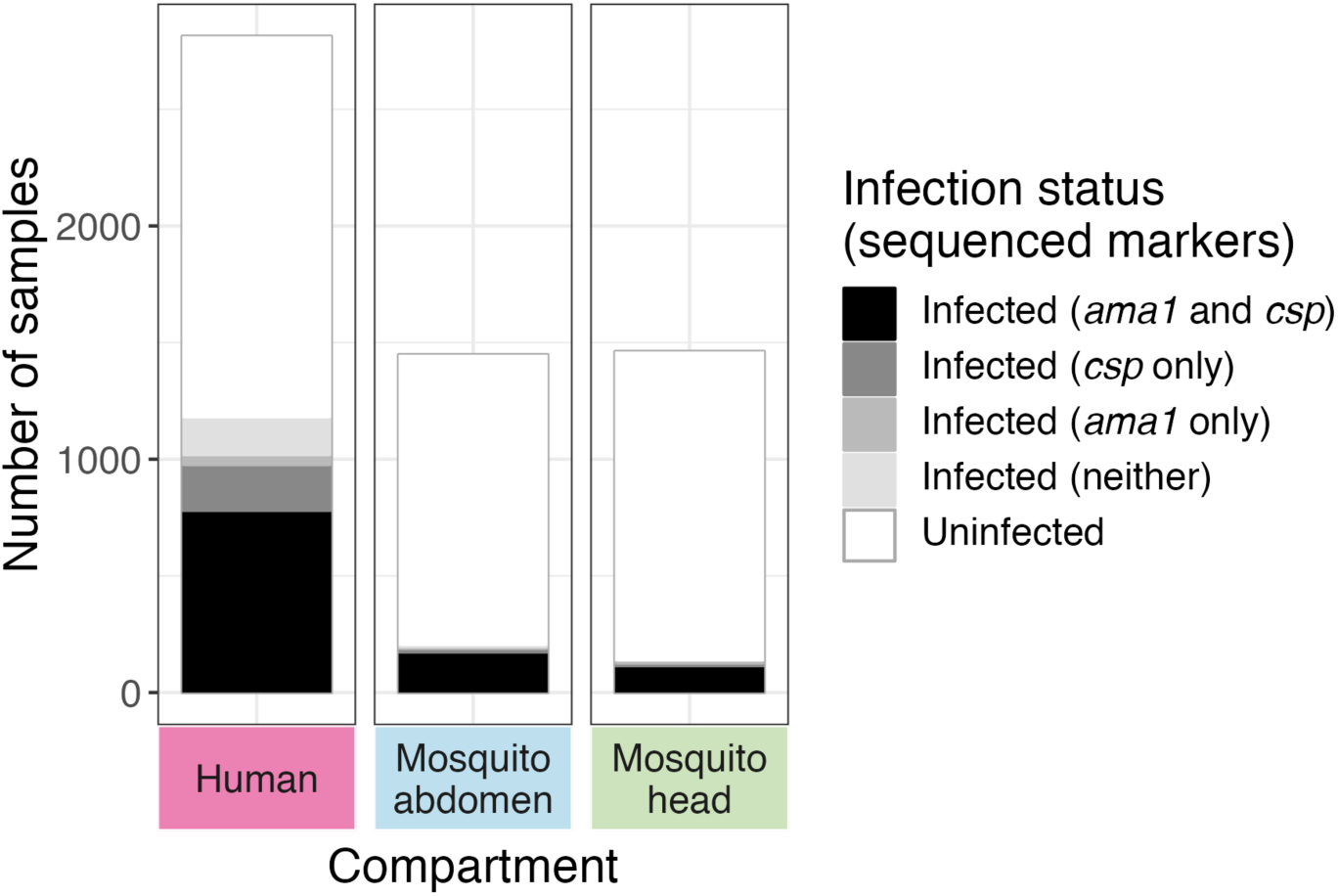
Overview of sample infection status. Samples with and without *P. falciparum* infections, including what markers were sequenced for each infected sample.

**Figure S2:**
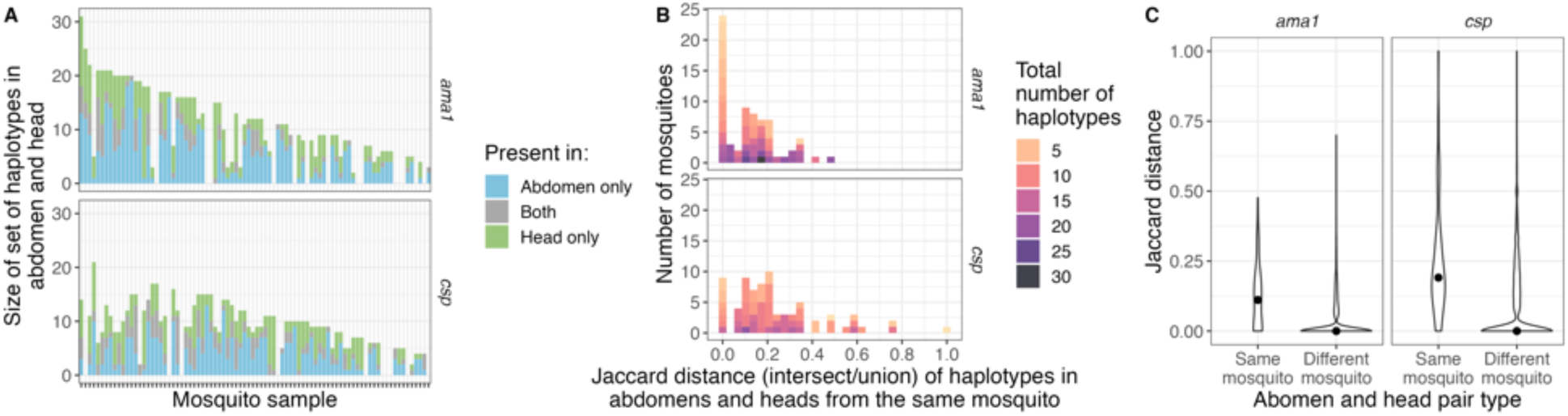
Summary of haplotypes found in the mosquito abdomen, mosquito head, or both compartments. (A) Sample-level counts. (B) Jaccard distance of haplotypes in abdomens and heads from the same mosquito, calculated as the intersect over the union of haplotypes in each mosquito. Color indicates the total number of haplotypes present in the mosquito, or the size of the union. (C) Jaccard distance of haplotypes observed in abdomens and heads compared between the same mosquito to that of different mosquitoes. The point is the median value.

**Figure S3:**
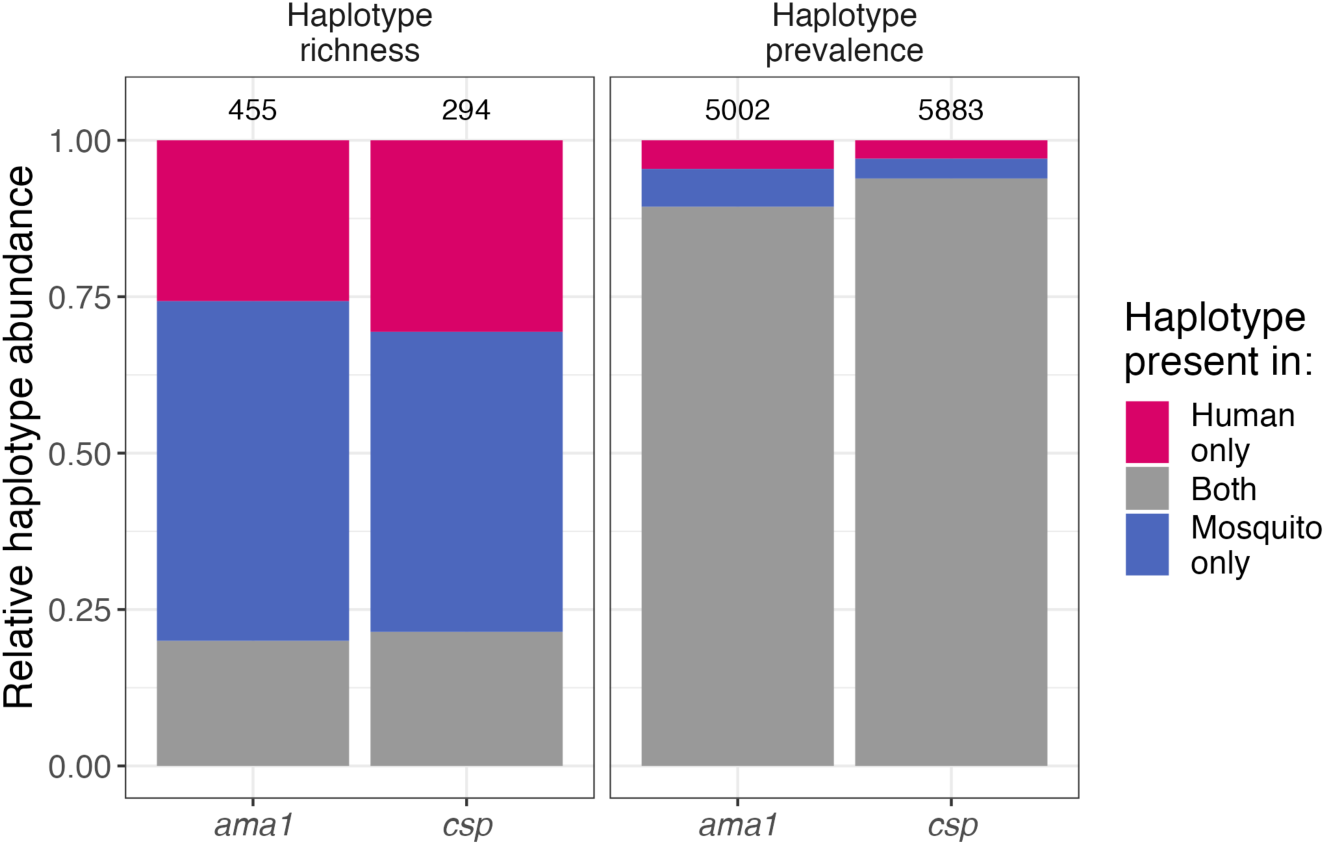
Haplotype richness and evenness across host compartments. Mean proportion of haplotypes present in each compartment, expressed as haplotype richness (left, including each unique haplotype once) and sample haplotype prevalence (right, counting the number of times each haplotype was observed).

**Figure S4:**
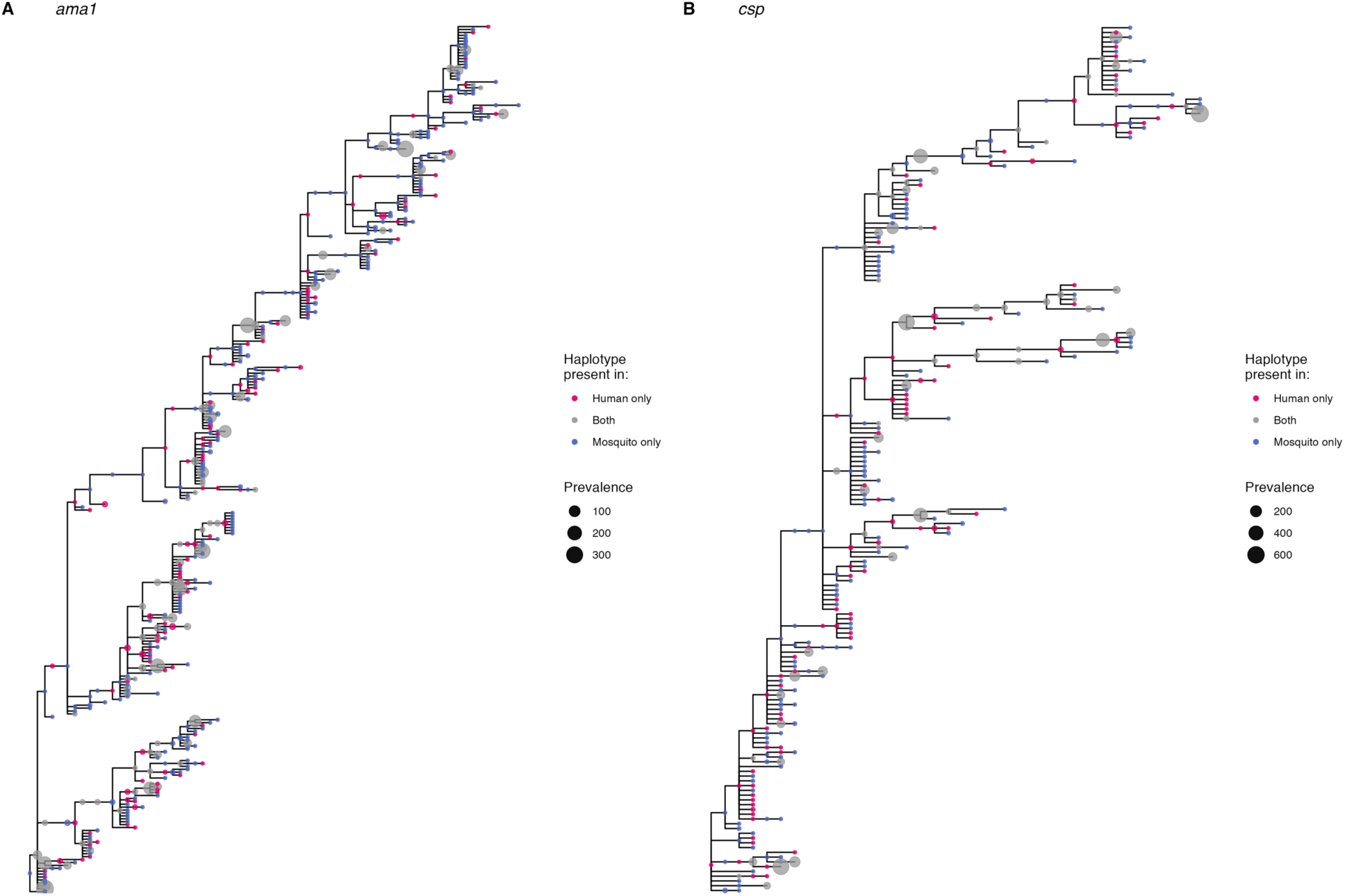
Randomized minimum spanning trees for nucleotide haplotype sequences.

**Figure S5:**
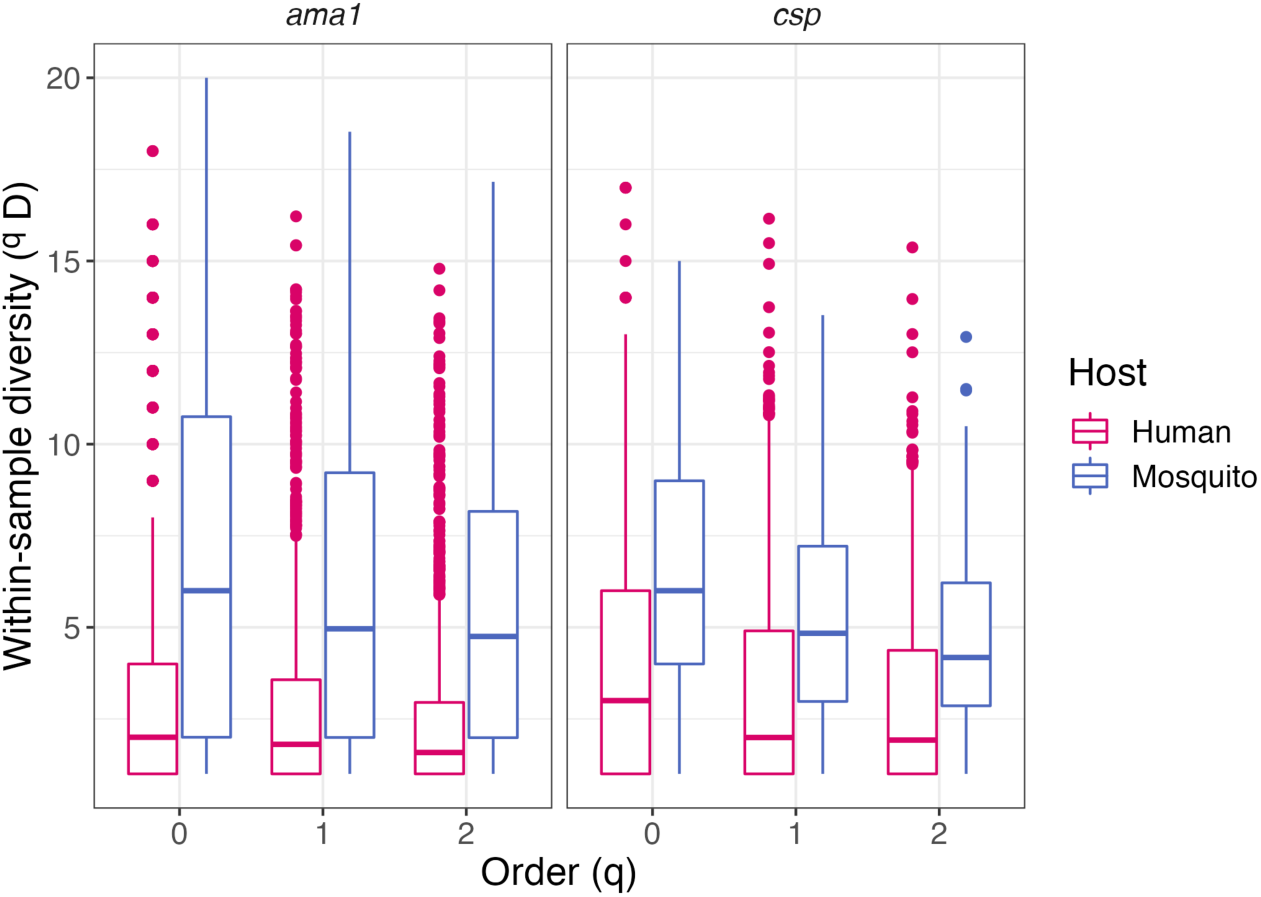
Within-sample diversity. When q = 0, the within-sample diversity is equivalent to the multiplicity of infection.

**Figure S6:**
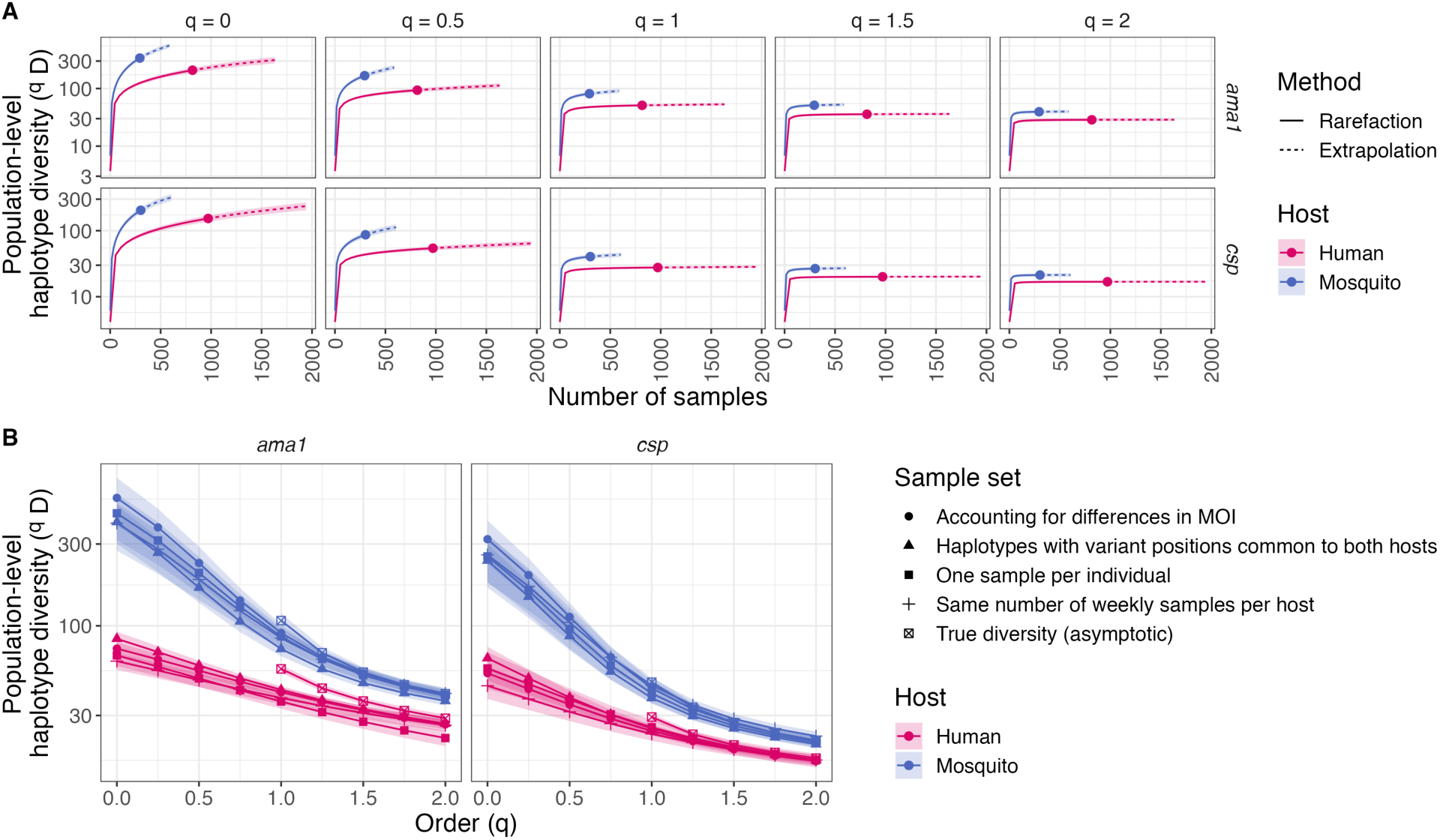
Mosquito haplotype populations are more diverse than human haplotype populations. (A) Rarefaction curves for various orders of diversity (*q*). True (asymptotic) diversity can be calculated after the rarefaction curve flattens out; otherwise, the computed true diversity is a lower bound. Comparisons can be made for true diversity at orders of diversity above q = 1 in our dataset because the human diversity curve flattens out and is lower than the mosquito diversity curve (which is a minimum bound on diversity). (B) Sensitivity analyses comparing diversity between humans and mosquitoes for subsampled data.

**Figure S7:**
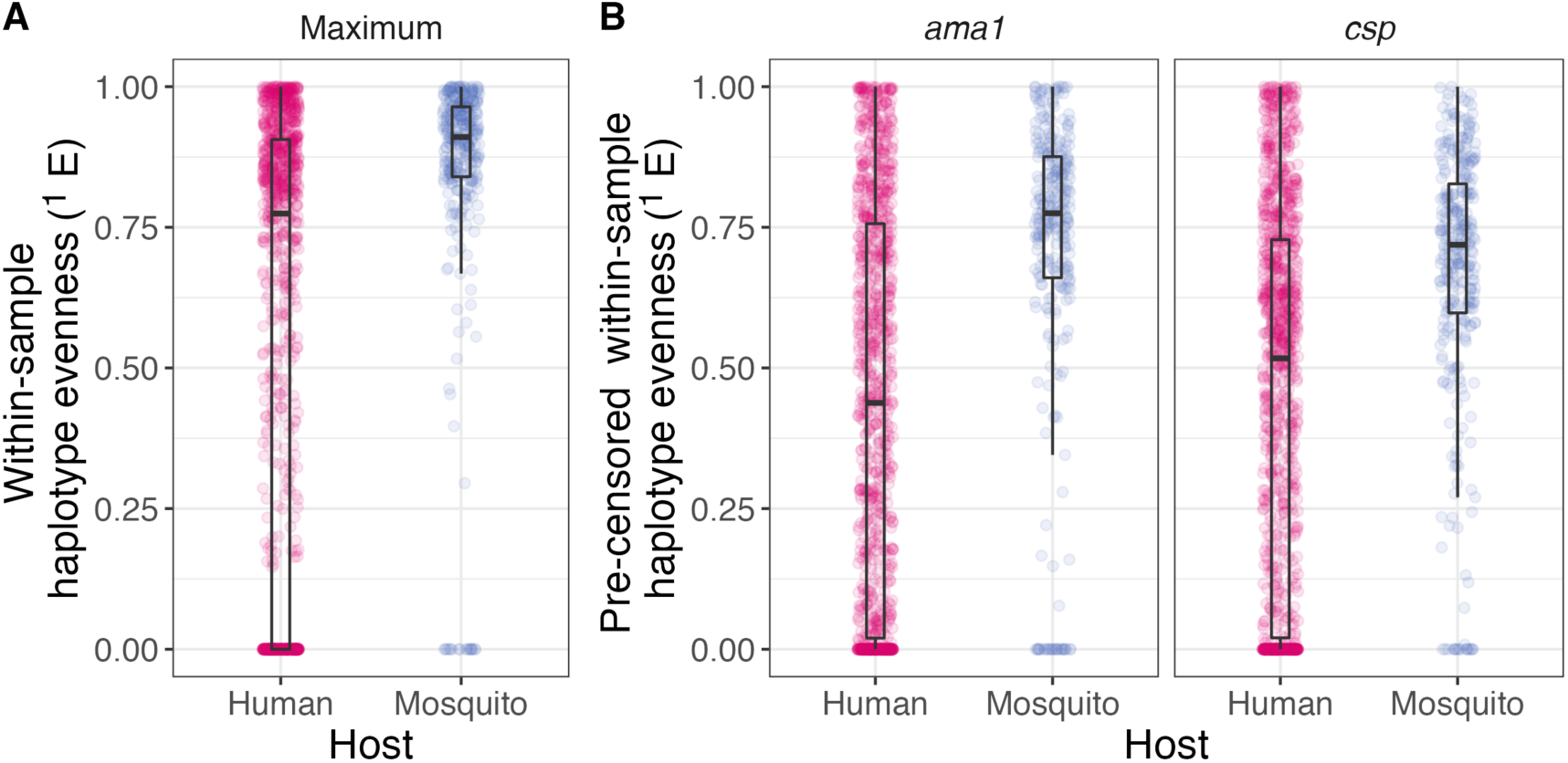
Haplotype evenness sensitivity analyses. (A) Taking the maximum evenness value between *ama1* and *csp*. (B) For pre-censored haplotype read counts. Both sensitivity analyses show the same trend as Figure 3B in the main text.

